# RBM43 links adipose inflammation and energy expenditure through translational regulation of PGC1α

**DOI:** 10.1101/2023.01.06.522985

**Authors:** Phillip A. Dumesic, Sarah E. Wilensky, Symanthika Bose, Jonathan G. Van Vranken, Steven P. Gygi, Bruce M. Spiegelman

## Abstract

Adipose thermogenesis involves specialized mitochondrial function that counteracts metabolic disease through dissipation of chemical energy as heat. However, inflammation present in obese adipose tissue can impair oxidative metabolism. Here, we show that PGC1α, a key governor of mitochondrial biogenesis and thermogenesis, is negatively regulated at the level of mRNA translation by the little-known RNA-binding protein RBM43. *Rbm43* is expressed selectively in white adipose depots that have low thermogenic potential, and is induced by inflammatory cytokines. RBM43 suppresses mitochondrial and thermogenic gene expression in a PGC1α-dependent manner and its loss protects cells from cytokine-induced mitochondrial impairment. In mice, adipocyte-selective *Rbm43* disruption increases PGC1α translation, resulting in mitochondrial biogenesis and adipose thermogenesis. These changes are accompanied by improvements in glucose homeostasis during diet-induced obesity that are independent of body weight. The action of RBM43 suggests a translational mechanism by which inflammatory signals associated with metabolic disease dampen mitochondrial function and thermogenesis.

## INTRODUCTION

Obesity is predicted to affect half of American adults—and over one billion adults worldwide—by 2030^1^. Many of these people will suffer from its metabolic sequelae such as diabetes and cardiovascular disease. In this context, the manipulation of adipose thermogenesis has garnered attention as a potential therapeutic strategy. As opposed to white fat cells, which function to store chemical energy, thermogenic adipocytes contain numerous mitochondria and achieve extraordinary levels of fuel oxidation, driving futile chemical cycles that expend chemical energy as heat. Thermogenic fat cells include brown as well as beige, a distinct cell type that arises within white fat upon stimulation^2^. Adipose thermogenesis is crucial for mammalian body temperature defense, but also confers metabolic benefits including increased energy expenditure, improved glucose and lipid homeostasis, and improved insulin sensitivity^3^. Active thermogenic fat is found in humans, though it remains uncertain whether its amount suffices as a cellular basis for therapy^4–6^. As such, a major goal is to understand means by which white fat can be made to take on a more thermogenic identity.

A fundamental challenge to the use of thermogenic fat for metabolic therapy is that obesity itself dampens thermogenesis. In humans, older, obese, or diabetic populations possess less active thermogenic fat^7,8^. In mice, obesity impedes thermogenic fat development and activity^9^. The low-grade inflammatory state that arises during obesity is thought to be a major cause. For instance, TNFα, IL-1β, interferon, and innate immune sensor signaling have all been implicated in suppression of thermogenesis^10–15^. The mechanisms by which inflammatory signals affect mitochondrial function are of great interest, as they might reveal means to preserve thermogenesis and normal mitochondrial function in the presence of inflammation, while also providing insight into the pathophysiology of obesity.

The transcriptional coactivator PGC1α was discovered by its ability to regulate thermogenic gene expression in brown fat^16^. It has since been established as a dominant regulator of mitochondrial biogenesis and oxidative metabolism in most if not all tissues^17^. By coordinating mitochondrial biogenesis with the expression of genes that carry out futile chemical cycles^16,18^, PGC1α is central to thermogenic fat function: its expression correlates with thermogenic potential across adipose depots, and its genetic disruption reduces thermogenesis and causes insulin resistance^19^. Though regulated via increased transcription by β-adrenergic signaling^20^, recent studies have highlighted the *Ppargc1a* mRNA itself as an important node of regulation^21–25^. For instance, our own work characterizing a mouse model of elevated *Ppargc1a* translation demonstrated that this mode of control is sufficient to increase oxidative metabolism in multiple tissues and to confer protection from ischemic kidney injury^26^. Here, our efforts to understand inputs into *Ppargc1a* translational control led to identification of RBM43, an RNA-binding protein induced by inflammatory cytokines that negatively regulates adipose thermogenesis via PGC1α.

## RESULTS

### The RNA-binding protein RBM43 is a negative post-transcriptional regulator of *Ppargc1a* and oxidative metabolism

To identify post-transcriptional regulators of oxidative metabolism, we studied RNA-binding proteins whose expression was coordinated with adipose thermogenesis, a process that involves extensive remodeling of mitochondrial content and function. Candidates of interest were: (1) RNA-binding proteins or members of a ribonucleoprotein complex^27^; (2) expressed preferentially in either white or thermogenic adipocytes^28^; and (3) regulated upon cold exposure in brown and inguinal white adipocytes^28^. After exclusion of protein products essential for cellular survival or localized to the mitochondria^29,30^, 25 candidates remained; we individually knocked these down in primary inguinal white adipose tissue (iWAT) adipocytes using siRNA (Figure S1A). We subsequently determined whether knockdown of each candidate affected the protein:mRNA ratio of PGC1α. The screen’s top hit, both in terms of protein:mRNA ratio change and its statistical significance, was *Rbm43* (Figure S1B). *Rbm43* encodes a 39 kDa polypeptide with a single N-terminal RNA recognition motif. It has been implicated in post-transcriptional repression of mRNAs involved in hepatocellular carcinoma G2/M progression^31^.

In follow-up experiments, two independent siRNAs targeting *Rbm43* similarly increased the PGC1α protein:mRNA ratio (1.9-fold and 1.9-fold) (Figures 1A,B). This effect was not accompanied by a change in the PGC1α protein half-life when measured during cycloheximide treatment (Figure 1C), suggesting an increase in *Ppargc1a* mRNA translation. The ability of *Rbm43* knockdown to increase PGC1α protein was not suppressed by the induction of *Ppargc1a* transcription using isoproterenol, consistent with the idea that RBM43 acts on *Ppargc1a* in a manner other than transcriptional control (Figure S1C).

**Figure 1.**
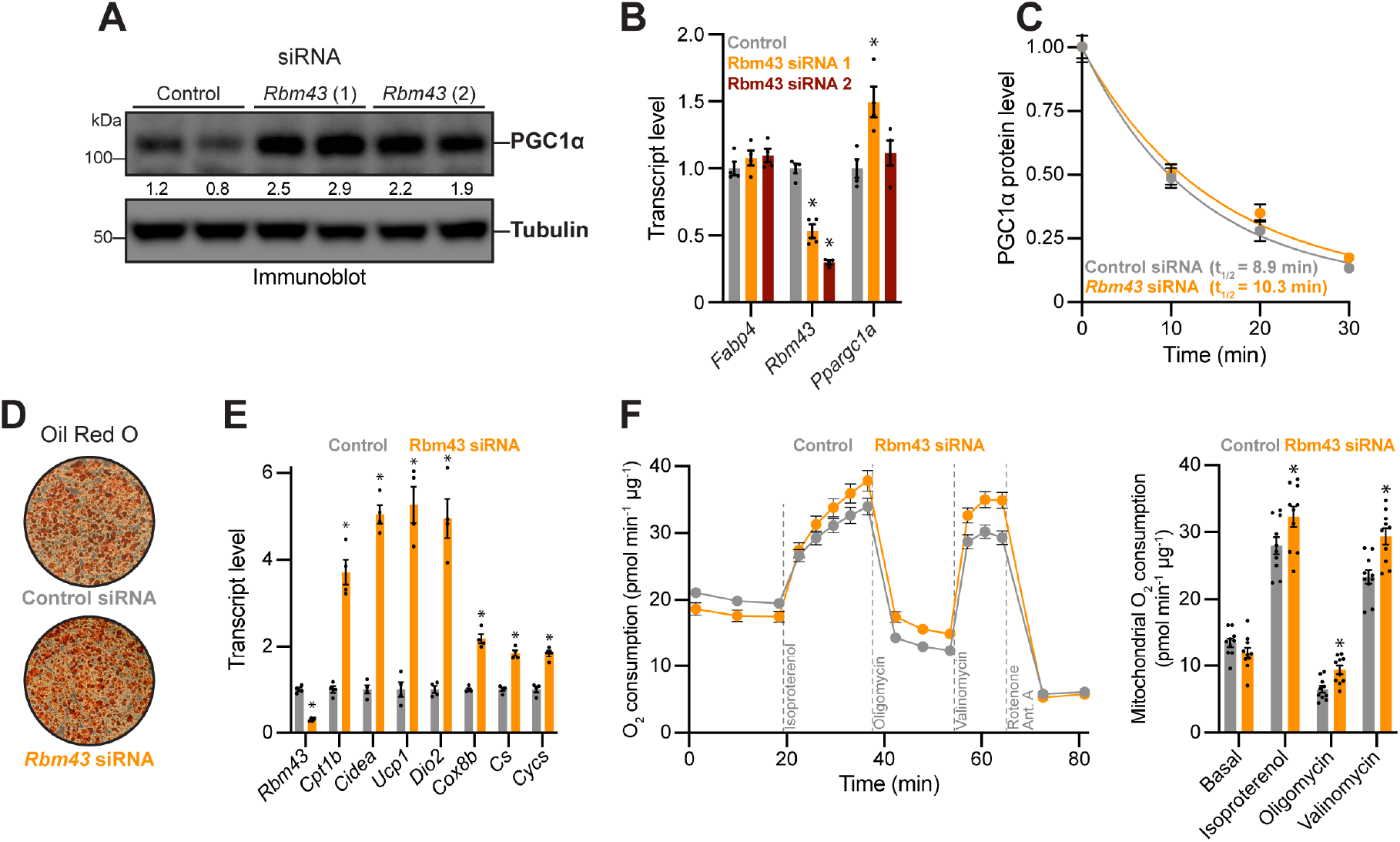
*Rbm43* knockdown elevates PGC1α protein output and mitochondrial respiration in primary adipocytes. **(A-B)** PGC1α protein (A) and mRNA (B) levels after knockdown of *Rbm43* in primary iWAT adipocytes. n=2 (protein) or 3 (mRNA). PGC1α immunoblot densitometry is normalized to signal in control samples. *p<0.01. **(C)** Half-life of PGC1α protein in *Rbm43* knockdown primary iWAT adipocytes, as measured using cycloheximide treatment. n=3. **(D)** Oil Red O staining of primary iWAT adipocytes after *Rbm43* knockdown. **(E)** RT-qPCR measuring mRNAs associated with thermogenesis and mitochondrial content in primary iWAT adipocytes after *Rbm43* knockdown. n=4.*p<0.001. **(F)** Total (left) and mitochondrial (right) oxygen consumption by primary iWAT adipocytes after *Rbm43* knockdown. Uncoupled respiration was measured in the presence of oligomycin; maximal respiration was measured in the presence of valinomycin. n=10. *p<0.05.

We next examined broader changes in the oxidative metabolism gene expression program, as well as respiration, in adipocytes depleted of *Rbm43*. Knockdown of *Rbm43* increased expression of genes involved in fatty acid oxidation, thermogenesis, and mitochondrial function without visibly affecting adipogenesis (Figures 1D,E). These changes were associated with elevated total and uncoupled mitochondrial respiration, as measured by oxygen consumption (Figure 1F). These data identify RBM43 as a negative post-transcriptional regulator of *Ppargc1a* and oxidative metabolism in adipocytes.

### RBM43 is selectively expressed in white fat and induced by inflammatory cytokines

The influence of RBM43 on oxidative metabolism led us to explore the regulation of its mRNA. In mice housed at 22 °C, *Rbm43* was detected in all mouse tissues examined, with highest levels in white adipose tissue (both inguinal and visceral) and spleen (Figure S2A). Lowest *Rbm43* levels were detected in classical brown adipose tissue, where they were 20% the level of *Rbm43* in white fat (Figures 2A,S2A). Assessment of *Rbm43* expression specifically in adipocytes using published translating ribosome affinity purification (TRAP) data confirmed its selectivity for white fat^28,32^ (Figure 2B). When mice were exposed to cold temperatures, *Rbm43* expression decreased in both WAT and BAT, including 4-6-fold decreases in adipocytes themselves (Figures 2C,D). Temperature-dependent regulation is a common feature of genes controlled by cold-induced adrenergic signaling. Indeed, primary iWAT adipocytes significantly decreased *Rbm43* expression in response to adrenergic agonists or agents that increase cellular cAMP (Figure 2E).

**Figure 2.**
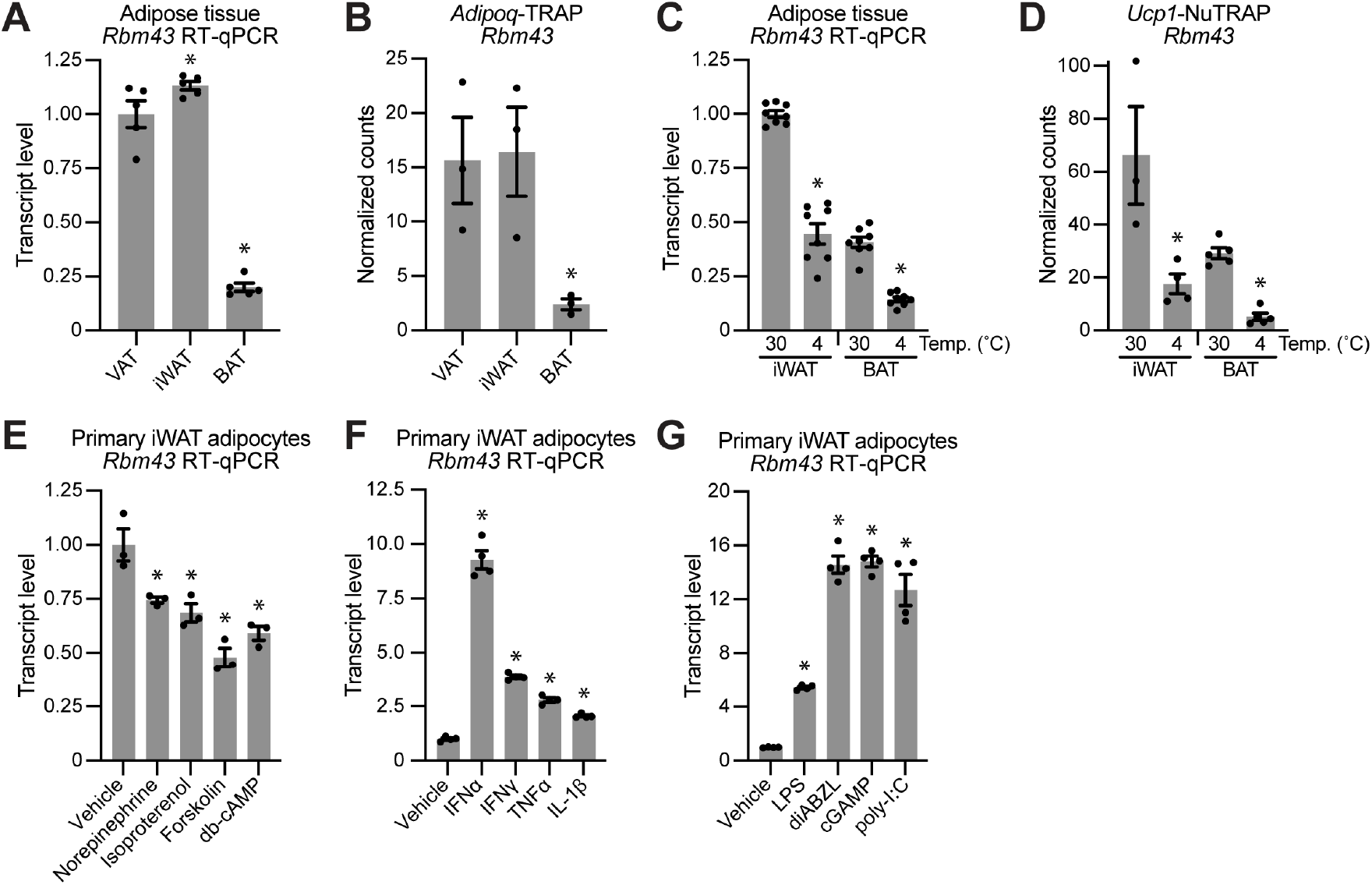
*Rbm43* is regulated by adrenergic signaling and inflammatory cytokines. **(A)** *Rbm43* expression in mouse adipose depots at 22°C. n=5. *p<0.05 vs VAT. **(B)** Adipocyte-specific expression of *Rbm43* in mouse adipose depots at 22°C, as assessed by *Adipoq-TRAP* (NCBI GEO: GSE103617). n=3. *p<0.05 vs VAT. **(C)** *Rbm43* expression in mouse adipose depots in response to cold exposure. n=8. *p<0.001 vs corresponding 30 °C. **(D)** Adipocyte-specific expression of *Rbm43* in mouse adipose depots in response to cold exposure, as assessed by *Ucp1-TRAP* (NCBI GEO: GSE108077). n=3-5. *p<0.05 vs corresponding 30 °C. **(E)** *Rbm43* expression in primary iWAT adipocytes after 3 h treatment with agents to activate adrenergic signaling. n=3. *p<0.005 vs vehicle control. **(F)** *Rbm43* expression in primary iWAT adipocytes after 4 h treatment with indicated cytokines. n=4. *p<0.005 vs vehicle control. **(G)** *Rbm43* expression in primary iWAT adipocytes after 8 h treatment with indicated agents to activate innate immune signaling. n=4. *p<0.001 vs vehicle control.

Examination of the promoter region of mouse *Rbm43* revealed potential binding sites for interferon regulatory factors (IRFs) and NF-κB, raising the possibility that cytokine signaling regulates *Rbm43^33^*. Primary iWAT adipocytes increased *Rbm43* expression upon treatment with type I or type II interferon, TNFα, or IL-1β, with the largest effect (9-fold) induced by IFNα (Figure 2F). The interferon response can be initiated by not only cell surface receptors, but also intracellular sensors of the innate immune system, such as Toll-like receptors, RIG-I, and cGAS-STING^34^. Treatment of primary iWAT adipocytes with lipopolysaccharide (a TLR4 ligand) and poly-inosine:cytosine (a ligand of TLR3 and activator of RIG-I/MDA5) both induced *Rbm43* (Figure 2G). Activators of cGAS-STING signaling (diABZL and cGAMP) had an even larger effect, increasing *Rbm43* mRNA 15-fold (Figure 2G). Together, these results indicate that *Rbm43* is subject to negative regulation by adrenergic signaling and positive regulation by pro-inflammatory cytokines as well as intracellular signaling of innate immune receptors.

### RBM43 acts through PGC1α to control oxidative metabolism gene expression

We next sought to determine the global gene expression program regulated by RBM43, and the extent to which this program operates through PGC1α. To this end, we knocked down *Rbm43* in *Ppargc1a^fl/fl^* adipocytes, in the presence of adenoviral *Cre* expression or *Gfp* control. Although >90% of adipocytes were transduced by adenovirus (as judged by adenoviral GFP), Cre disrupted only ~50% of *Ppargc1a^fl^* alleles (Figure S3A). This provided a scenario in which PGC1α protein was reverted to near-baseline levels during *Rbm43* knockdown, thereby allowing precise experimental isolation of the consequences of RBM43-dependent PGC1α regulation.

Upon knockdown of *Rbm43*, mRNAs corresponding to 693 genes were elevated (log_2_FC>0.3; p_adj_<0.005) (Figure 3A). Oxidative phosphorylation, adipogenesis, and fatty acid metabolism were the three most overrepresented Hallmark gene sets among elevated genes. These gene sets contain central components of oxidative phosphorylation metabolism, including many direct and indirect targets of PGC1α (Figure 3B). Gene set enrichment analysis confirmed strong induction of a previously-defined signature of PGC1α activity^26^ (Figure 3D). All of these results suggest that PGC1α activity is elevated upon RBM43 loss. Indeed, Cre-mediated reduction of PGC1α reversed most of the upregulated genes: 492 (71%) reverted toward their baseline values (as determined by a Z>0.67 change) (Figure 3A).

**Figure 3.**
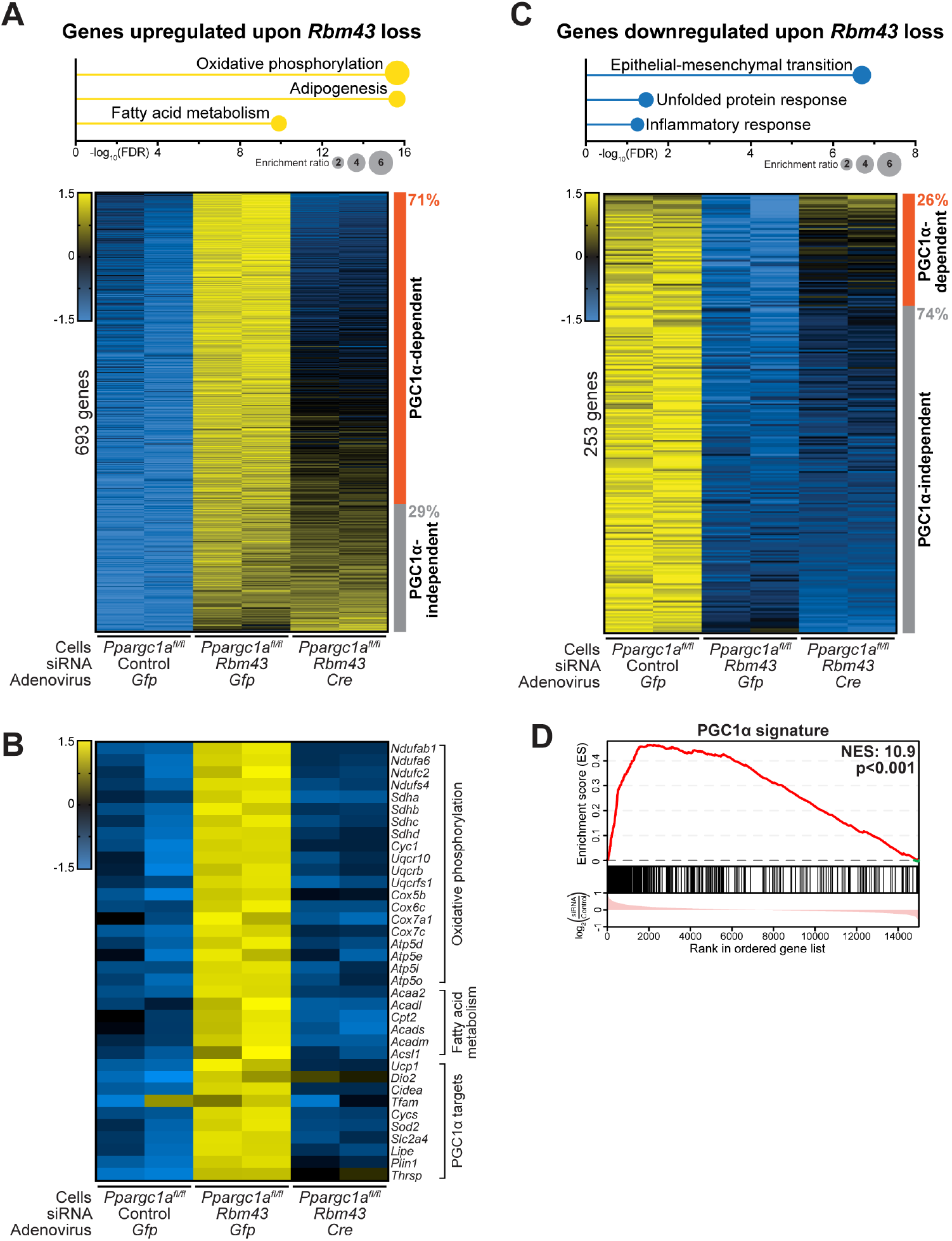
RBM43 acts through PGC1α to control the adipocyte oxidative phosphorylation gene expression program. **(A)** *Top:* Top three Hallmark gene sets overrepresented amongst the genes upregulated upon *Rbm43* knockdown. *Bottom:* RNA-seq Z-score heatmap of the 693 genes upregulated upon *Rbm43* knockdown (log_2_FC>0.3; p_adj_<0.005). The fraction of these genes whose expression change is reversed upon *Ppargc1a* knockout (Z-score reversion >0.67) is shown. n=2. **(B)** RNA-seq Z-score heatmap of representative genes involved in oxidative phosphorylation, fatty acid metabolism, and direct PGC1α targets. n=2. **(C)** *Top:* Top three Hallmark gene sets overrepresented amongst the genes downregulated upon *Rbm43* knockdown. *Bottom:* RNA-seq Z-score heatmap of the 253 genes downregulated upon *Rbm43* knockdown (log_2_FC>0.3; p_adj_<0.005). The fraction of these genes whose expression change is reversed upon *Ppargc1a* knockout (Z-score reversion >0.67) is shown. n=2. **(D)** Gene set enrichment plot assessing enrichment of PGC1α signature genes amongst genes altered by *Rbm43* knockdown.

We next examined the 253 genes that were reduced upon *Rbm43* knockdown (Figure 3C). Epithelial-mesenchymal transition, unfolded protein response, and inflammatory response were the most overrepresented gene sets. All of these programs have been associated with mitochondrial dysfunction and obesity^35–39^, raising the question of whether their decrease upon *Rbm43* knockdown is a direct consequence of RBM43 loss, or rather a secondary effect of PGC1α and its influence on mitochondria. In support of the former, Cre-mediated reduction of PGC1α reverted only 66 (26%) of the genes downregulated upon RBM43 loss (Figure 3C).

Thus, these results delineate two independent arms of the gene expression program controlled by RBM43. In one arm, RBM43 inhibits genes related to oxidative metabolism, mainly via its suppression of PGC1α. In the second arm, RBM43 activates genes related to cellular identity, stress, and inflammation in a largely PGC1α-independent manner.

### Adipocyte-selective *Rbm43* knockout increases PGC1α translation, mitochondrial biogenesis, and adipose thermogenesis in mice

We generated a conditional mouse model of *Rbm43* loss of function, in which exon 3 was flanked by *loxP* sites. *Adiponectin-Cre* was then used to disrupt *Rbm43^fl/fl^* in an adipocyte-selective manner (*Rbm43-AKO)^40^. Rbm43-AKO* mice were born at expected Mendelian ratios with no gross abnormalities. In young adult mice fed a normal chow diet, the weights of brown and white fat depots did not differ between *Rbm43-AKO* mice and their *Rbm43^fl/fl^* littermates, though the *Rbm43-AKO* inguinal WAT depots took on a browner color suggestive of increased thermogenic fat content (Figures S4A,B). Isolation of iWAT adipocytes by flotation confirmed >85% disruption of the *Rbm43* allele, as well as elevations in genes preferentially expressed in thermogenic fat, such as *Ucp1, Cox8b*, and *Elovl3* (Figure S4C).

Isobaric peptide labeling using TMTpro reagents enabled quantitative determination of RBM43’s effect on the global proteome of isolated iWAT adipocytes^41^. The Hallmark gene sets most upregulated in *Rbm43-AKO* cells were oxidative phosphorylation, adipogenesis, and fatty acid metabolism, similar to the results obtained upon siRNA-mediated *Rbm43* knockdown *in vitro* (Figures 4A-C). Across all annotated oxidative phosphorylation complex components^42^, mean protein increase in *Rbm43-AKO* cells was 15%, a result confirmed by immunoblot (Figures 4D-F). In addition, the PGC1α gene expression signature was strongly induced, consistent with an elevation in PGC1α action (Figure 4G). Of note, the PGC1α-independent effects of RBM43 defined *in vitro* were again observed upon *in vivo* loss of RBM43: downregulated gene sets included epithelial-mesenchymal transition and inflammatory response (Figures 4B,S4D). Thus, the multiple RBM43-dependent programs defined *in vitro* were recapitulated upon *in vivo* loss of RBM43 in adipocytes.

**Figure 4.**
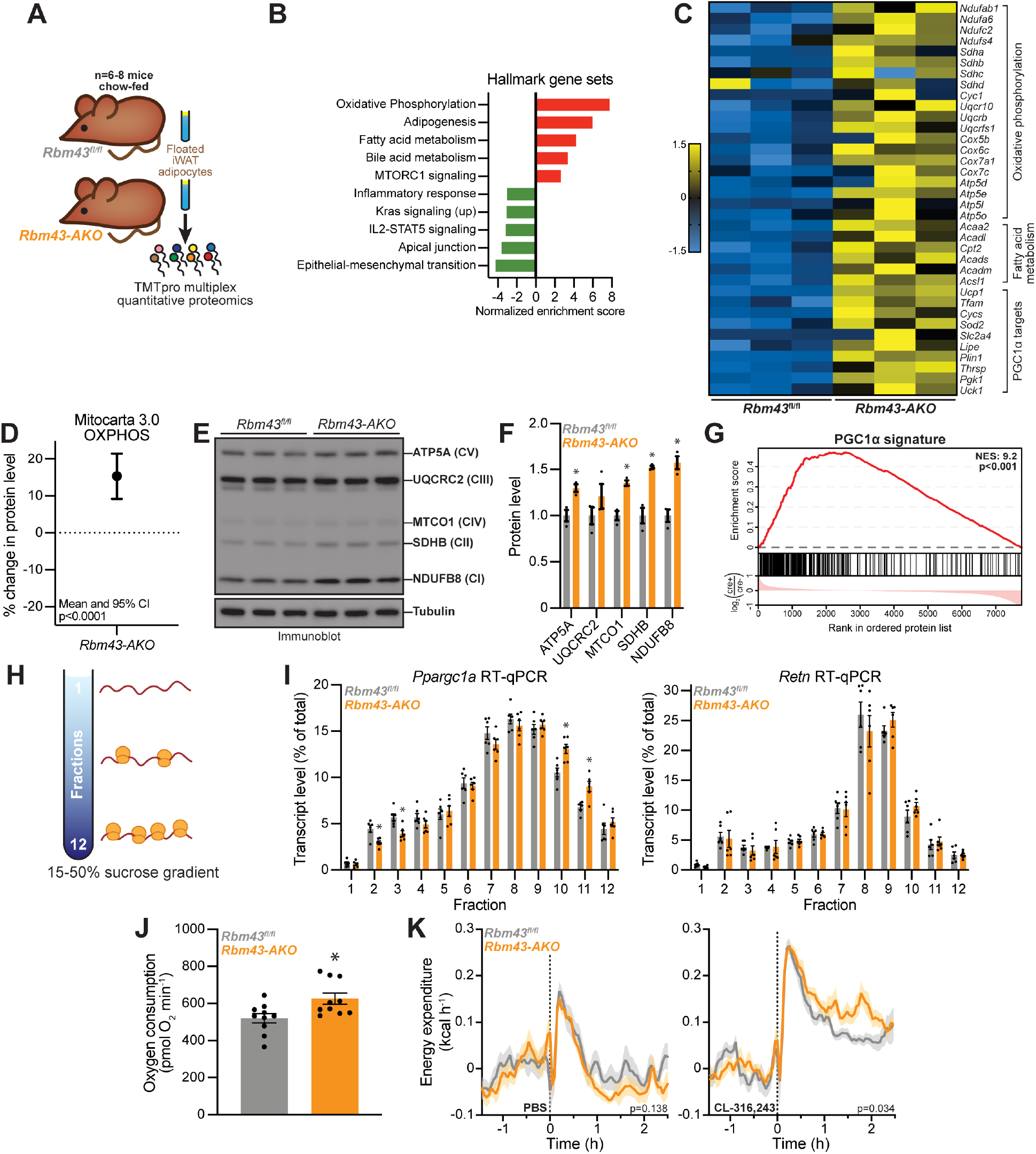
PGC1α translation, mitochondrial biogenesis, and thermogenesis are elevated upon RBM43 loss in fat. **(A)** Strategy for *Rbm43-AKO* iWAT proteomics. **(B)** Top 5 Hallmark gene sets enriched in positive or negative direction in *Rbm43-AKO* proteomic data, as assessed by gene set enrichment analysis. **(C)** Proteomics Z-score heatmap of representative proteins involved in oxidative phosphorylation, fatty acid metabolism, and direct PGC1α target genes. n=3. **(D)** Percent protein abundance change (*Rbm43-AKO* vs *Rbm43^fl/fl^*) for all proteins annotated as oxidative phosphorylation complex components (Mitocarta 3.0). Mean and 95% confidence interval are shown. n=128. **(E-F)** Immunoblot (E) and corresponding densitometry (F) of oxidative phosphorylation complex components in iWAT adipocytes from *Rbm43^fl/fl^* and *Rbm43-AKO* mice. n=3. *p<0.05. **(G)** Gene set enrichment plot assessing enrichment of PGC1α signature gene set among proteins upregulated in *Rbm43-AKO* iWAT proteomics data. **(H)** Schematic of polysome profiling experiment, in which sucrose density gradients resolve highly translated mRNAs (dense fractions) from lowly translated mRNAs (light fractions). **(I)** Profiles of *Ppargc1a* (left) and *Retn* (right) on polysomes isolated from iWAT adipocytes of *Rbm43^fl/fl^* and *Rbm43-AKO* mice. n=6. *p<0.005. **(J)** Respiration in minced iWAT tissue (5 mg), as measured by oxygen consumption in Seahorse Bioanalyzer. n=10. *p<0.05. **(K)** Energy expenditure in *Rbm43^fl/fl^* and *Rbm43-AKO* mice after administration of PBS (left) or CL-316,243 (right). n=10-14.

The strong increase in PGC1α target gene expression in the absence of elevated *Ppargc1a* mRNA was suggestive of post-transcriptional control. We therefore assessed active translation of *Ppargc1a* in adipocytes isolated from *Rbm43-AKO* mice. Floated iWAT adipocytes were treated with cycloheximide to preserve ribosome-mRNA interactions, followed by lysis and sucrose density gradient centrifugation to resolve lowly-translated mRNAs (low density fractions: free mRNA and monosomes) from highly-translated mRNAs (high density fractions: polysomes)^43^ (Figure 4H). The overall ribosome distribution in the gradient, as judged by immunoblot of RPS3, was similar between *Rbm43-AKO* and Rbm43^fl/fl^ cells (Figure S4E). The *Ppargc1a* mRNA, however, was shifted to higher-density fractions in the context of RBM43 loss (Figure 4I). The distribution of the *Retn* mRNA, which served as a control, was not altered. These results suggest that an increase in *Ppargc1a* translation caused by loss of RBM43 contributes to the elevation of oxidative metabolism gene expression seen in adipose tissue of *Rbm43-AKO* mice.

We subsequently tested whether the gene expression changes caused by loss of RBM43 in adipocytes were accompanied by an increased capacity for oxidative metabolism. Minced iWAT was capable of elevated respiration in the context of RBM43 loss, as measured by oxygen consumption in a Seahorse bioanalyzer (Figure 4J). Next, to assess the adipocyte compartment specifically, we treated living mice with the β3-adrenergic receptor agonist CL-316,243, followed by measurement of energy expenditure in metabolic cages. Whereas PBS control injections caused an equivalent stress response in both *Rbm43-AKO* mice and their *Rbm43^fl/fl^* littermates, CL-316,243 administration elicited significantly greater energy expenditure in the *Rbm43-AKO* mice (Figure 4K). These results indicate a greater capacity for thermogenesis in adipocytes lacking RBM43.

### RBM43 loss in adipocytes improves glucose tolerance independent of weight in HFD-fed mice

Given the potential for RBM43 to link cytokine signaling with mitochondrial regulation, we aimed to test whether its loss would protect adipocytes from the adverse effects of inflammation. First, we utilized a model of cytokine action on primary iWAT adipocytes in which 24 h treatment with TNFα broadly reduces expression of genes encoding mitochondrial proteins, including members of the TCA cycle (Figure S5A). Knockdown of *Rbm43* not only increased expression of multiple TCA cycle enzymes, but also protected against the TNFα-induced reduction in these genes. These results suggest that RBM43 is responsible, at least in part, for the effects of inflammatory cytokines on the expression of mitochondrial genes in fat.

To test the influence of inflammatory cytokines on adipose function *in vivo*, we examined glucose homeostasis in the context of high-fat diet (HFD) feeding, which induces obesity and adipose inflammation. In male mice fed a chow diet, neither body composition nor body weight differed between *Rbm43^fl/fl^* and *Rbm43-AKO* littermates (Figures 5A,B). Glucose homeostasis was largely unaltered in *Rbm43-AKO* mice as measured by intraperitoneal glucose tolerance test, with blood glucose levels significantly different between the genotypes only at the 30 min timepoint (Figure 5C). In female mice fed a high-fat diet for 12 weeks, neither body composition nor body weight differed between *Rbm43^fl/fl^* and *Rbm43-AKO* littermates (Figures 5D,E). Nevertheless, these *Rbm43-AKO* mice exhibited a substantial improvement in glucose tolerance (Figure 5F), indicating that mice lacking RBM43 in adipocytes are partially protected from the metabolic consequences of obesity.

**Figure 5.**
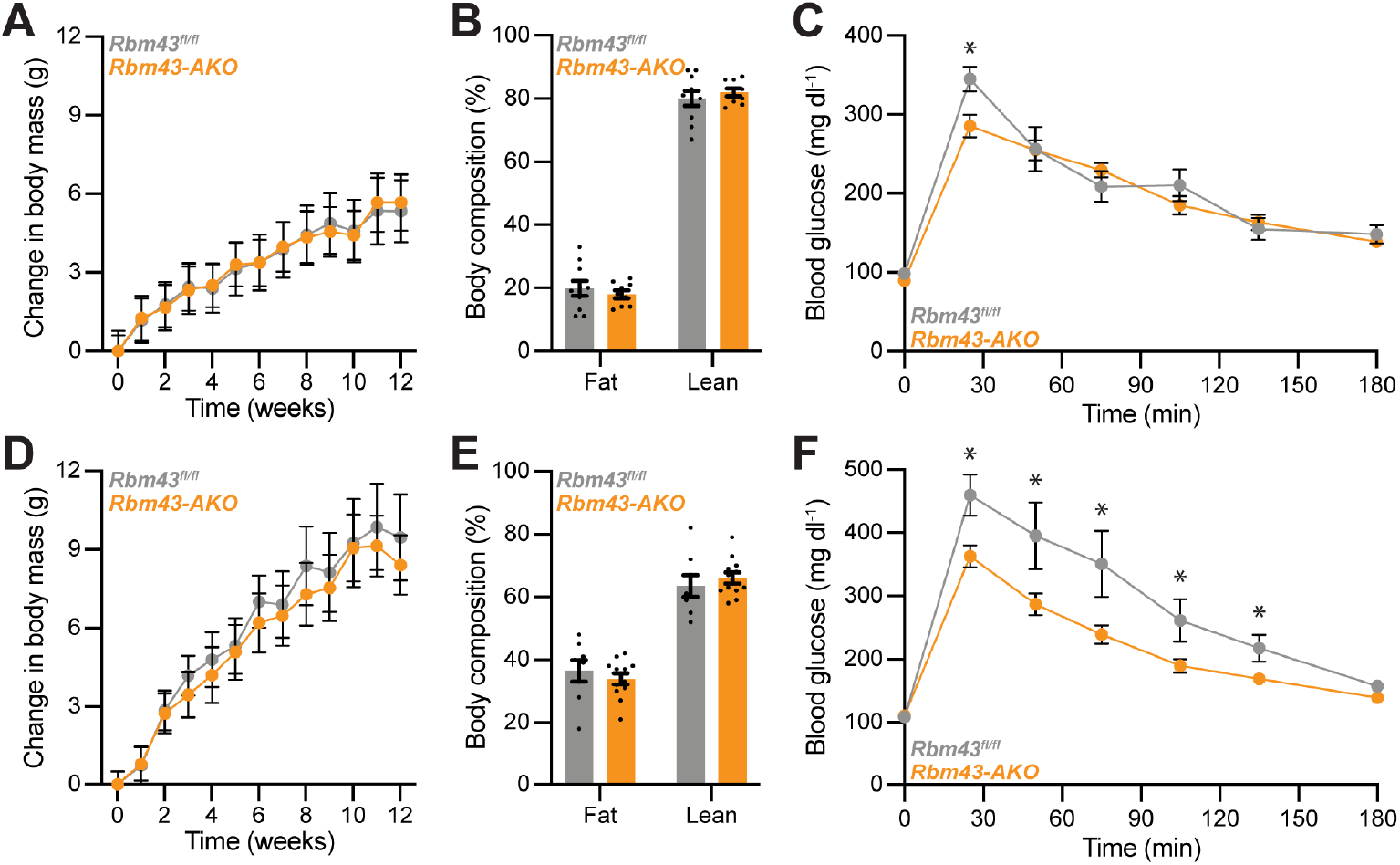
RBM43 loss in adipocytes improves glucose tolerance independent of body weight in mice fed a high-fat diet. **(A)** Body mass change during chow diet feeding to male *Rbm43^fl/fl^* and *Rbm43-AKO* mice at 22 °C. Total body weight did not significantly differ between genotypes. n=9-10. **(B)** Body composition of chow-fed *Rbm43^fl/fl^* and *Rbm43-AKO* male mice. n=9-10. **(C)** IP-GTT in chow-fed *Rbm43^fl/fl^* and *Rbm43-AKO* male mice. n=9-10. *p<0.05. **(D)** Body mass change during high fat diet feeding to female *Rbm43^fl/fl^* and *Rbm43-AKO* mice at 22 °C. Total body weight did not significantly differ between genotypes. n=8-12. **(E)** Body composition of high fat fed *Rbm43^fl/fl^* and *Rbm43-AKO* female mice after 11 weeks of diet. n=8-12. **(F)** IP-GTT in high fat fed *Rbm43^fl/fl^* and *Rbm43-AKO* female mice after 11 weeks of diet. n=8-12. *p<0.05.

## DISCUSSION

The salutary metabolic effects of adipose thermogenesis have stimulated interest in harnessing this physiology for the treatment of metabolic diseases. But these efforts are challenged by the intricate regulation governing the number and activity of thermogenic fat cells. Thermogenic adipocytes are generally outnumbered by energy-storing white adipocytes; harnessing the latent potential of white adipose tissue will require an understanding of how the thermogenic program is rendered absent or kept inactive in this cell type^44^. Furthermore, the pathophysiology of obesity actively impedes thermogenesis, in part through proinflammatory signaling, though the means by which these signals influence mitochondrial function remain unclear. Our central findings regarding the role of RBM43 in adipose tissue provide new insight into these questions.

First, we found that *Rbm43* is regulated in a manner opposite that of the thermogenesis program: it is expressed preferentially in white versus brown fat; suppressed by cold exposure and adrenergic signaling; and induced by cytokines and innate immune sensors. Much of this regulation likely occurs at the *Rbm43* promoter, which contains potential binding sites for interferon regulatory factors and NF-κB^33^. Interestingly, ChIP experiments demonstrate that PRDM16 also binds in this region^14^. PRDM16 is a critical positive regulator of the thermogenic fat transcriptional identity^45^, but it can also act as a repressor when bound to interferon-stimulated genes^14^. Consistent with this model, forced expression of PRDM16 reduces *Rbm43* expression in adipocytes^46^, whereas forced expression of IRF3 increases *Rbm43*^12^. This regulatory network suggests that RBM43 may contribute to the effects of IRF3, whose elevated expression in fat during obesity has been shown to suppress thermogenesis and insulin sensitivity^12,13^.

Second, we defined a major function of RBM43: to suppress the translation of *Ppargc1a* mRNA. Knockdown of *Rbm43* led to an increased PGC1α protein:mRNA ratio, without a change in PGC1α protein half-life. Protein:mRNA ratio is an imperfect measure for translational control because translation and mRNA stability are inherently coupled^47^, and its interpretation is complicated further for proteins like PGC1α that activate their own transcription^48^. Nevertheless, translational regulation is likely a significant part of RBM43’s action, since *Ppargc1a* mRNA, when assessed in polysome profiles, shifted to denser ribosomal fractions upon RBM43 loss. It remains to be determined which regions of the *Ppargc1a* mRNA are necessary for this regulation, though our experiments in primary adipocytes demonstrate that an intact uORF start codon is not required for RBM43’s effects. RBM43 does not appear to act through the previously described *Ppargc1a* uORF^26^.

Further analysis of RBM43’s genome-wide effects using RNA-seq epistasis analysis confirmed the centrality of PGC1α: of the 693 genes repressed by RBM43, >70% are controlled through PGC1α. In addition, a smaller, PGC1α-independent program is regulated (positively) by RBM43, which included genes involved in inflammation. These gene expression patterns were recapitulated in iWAT when *Rbm43* loss of function was assessed *in vivo* using our *Rbm43-AKO* mouse model. Nevertheless, it remains to be tested whether RBM43 regulates comparable genes across multiple cell types. Given that a post-transcriptional regulator requires its target mRNA in order to act, it is likely that RBM43’s actions in other cell types (or adipose depots) that express low levels of *Ppargc1a* will be less focused on mitochondrial biogenesis. Consistent with this idea, a recent report implicated RBM43 in the inhibition of genes involved in G2/M transition via post-transcriptional repression of *Ccnb1* (which encodes Cyclin B1, a protein lowly expressed in post-mitotic adipocytes)^31^.

Third, we demonstrated the thermogenic and metabolic impact of RBM43 loss in fat: substantial activation of PGC1α target genes, leading to elevated mitochondrial biogenesis, iWAT respiration, and CL-316,243-induced energy expenditure. Interestingly, this cellular phenotype did not lead to changes in weight gain or body composition on a high-fat diet, at least under the conditions and experimental duration described here. *Rbm43-AKO* mice did, however, show substantially improved glucose handling. This phenotype may represent a result complementary to that of fat-selective *Ppargc1a* knockout mice, which show decreased iWAT mitochondrial content and worsened insulin sensitivity on a HFD, independent of weight^19^. It also echoes the phenotype of visceral fat-selective deletion of *Zfp423* (a transcriptional repressor of the thermogenic identity), which shows improved insulin sensitivity on a HFD independent of weight^49^. These may represent examples of thermogenic fat contributing metabolic benefits in addition to energy expenditure *per se*. For instance, metabolic protection from HFD may arise in a cell-autonomous manner owing to the thermogenic program’s ability to influence mitochondrial biogenesis and function^50,51^. Alternatively, thermogenic fat cells may act non-cell autonomously to remodel the adipose milieu in ways that limit immunocyte recruitment, reduce fibrosis, and promote insulin sensitivity^3,52^. Thermogenic fat cells can also engage in inter-organ signaling that improves glucose handling in distant organs^53^. In this regard, it is interesting to note that a SNP near the human *RBM43* gene has been associated with type 2 diabetes in an African-American population^54^.

Finally, we note that there are likely circumstances in which the ability of RBM43 to link inflammatory cues to mitochondrial suppression is adaptive. For instance, interferon signaling aids the clearance of intracellular pathogens by promoting synthesis of antimicrobial molecules such as NO. The synthesis of these molecules can require mitochondrial suppression, which could conceivably be aided by RBM43^55^. Nevertheless, our results in the setting of overnutrition and metabolic inflammation highlight RBM43 as a potential mediator of maladaptive mitochondrial suppression.

## ACKNOWLEDGMENTS

This work was supported by National Institute of Health grants R01 DK119117 to B.M.S., K99 DK125722 to P.A.D.; JPB Foundation grant 6293803 to B.M.S.; and Damon Runyon Cancer Research Foundation Fellowship 120-17 to P.A.D. We thank the BIDMC Transgenic Mouse Core and the DFCI Molecular Biology Core Facilities for experimental assistance, and members of the Spiegelman laboratory for helpful discussions.

## AUTHOR CONTRIBUTIONS

P.A.D.: Conceptualization, investigation, formal analysis, writing, funding. S.E.W. and S.B.: investigation, data curation. J.G.V.: investigation, data curation. S.P.G.: Supervision, resources. B.M.S.: Supervision, conceptualization, writing, resources, funding.

## METHODS

### Mouse strains and husbandry

Mouse husbandry and experimentation were performed according to protocols approved by the Institutional Animal Care and Use Committee of the Beth Israel Deaconess Medical Center. Unless otherwise noted, mice were housed at 22°C under a 12 h light-dark cycle, with free access to food and water. A conditional allele of *Rbm43* was generated using the EASI-CRISPR method to flank exon 3 with *loxP* sites^56^. Fertilized eggs of C57BL/6J mice were microinjected with recombinant Cas9 (PNA Bio) and guide RNAs (Synthego) at the Beth Israel Deaconess Transgenic Core. Resulting pups were crossed to C57BL/6J mice and individual *Rbm43* alleles in the progeny were verified by Sanger sequencing as previously described^56^. Thereafter, genotyping was performed using PCR primers (TGATACTATTATTTGGGCCTTCC and CTTTGACCTTCCACGCTGAT), which distinguish a floxed allele (271 bp product) from a wild-type allele (237 bp product). Adipocyte-selective knockout of *Rbm43* was achieved using *Adiponectin-Cre* mice (JAX stock #028020)^40^.

### Primary adipocyte isolation and treatment

The stromal vascular fraction of iWAT was isolated from male and female C57BL/6J mice aged 4-10 weeks. Inguinal WAT was dissected and washed in Hank’s Buffered Saline Solution (HBSS) without calcium or magnesium (Corning). It was then minced and digested in HBSS containing 10 mg ml^-1^ Collagenase D (Roche), 3 U ml^-1^ Dispase II (Roche), and 10 mM CaCl_2_ for 45 min at 37 °C. After digestion, the cell suspension was combined with adipocyte culture medium (DMEM/F12 supplemented with 2.5 mM L-Alanyl-L-Glutamine, 10% fetal bovine serum [GeminiBio], 100 U ml^-1^ Penicillin [Gibco], 100 ug ml^-1^ Streptomycin [Gibco], and 0.1 mg ml^-1^ Primocin [Invivogen]) and filtered through a 100 μm cell strainer. Cells of the stromal vascular fraction were pelleted by centrifugation at 600 g for 5 min, after which they were resuspended in adipocyte culture medium, filtered through a 40 μm cell strainer, pelleted as above, and resuspended in adipocyte culture medium. The cells were plated on tissue culture plates and cultured at 37 °C with 10% CO2. Primocin was removed from the culture medium after 4 days.

Preadipocytes were cultured until 2 d after reaching confluency, then differentiated by addition of fresh adipocyte culture medium supplemented with 1 μM rosiglitazone (Cayman Chemical), 0.5 mM isobutylmethylxanthine (Sigma), 1 μM dexamethasone (Sigma), and 870 nM insulin (Sigma). Two days later (‘day 2’), and every 2 d thereafter, medium was replaced with fresh adipocyte culture medium supplemented with 1 μM rosiglitazone and 870 nM insulin. After day 7, rosiglitazone and insulin were omitted from the medium since differentiation was complete. Depending on assay, experiments were performed at day 8-10 of differentiation as noted below.

For *Ppargc1a* knockout experiments, primary adipocytes were prepared from *Ppargc1a^fl/fl^* mice (JAX #009666)^57^ and transfected with siRNA as described above. In addition, cells were transduced on day 1 of differentiation at a multiplicity of infection of 300 with purified adenovirus encoding *Gfp* or *Cre* (Boston Children’s Hospital Viral Core).

For experiments to test pathways that regulate *Rbm43* expression, primary adipocytes were prepared as described above, and treated on day 8 or 9 with the indicated agents: norepinephrine (2 μM, Sigma), isoproterenol (1 μM, Sigma), forskolin (10 μM, Sigma), db-cAMP (500 μM, Sigma), IFNα (25 ng ml^-1^, R&D Systems), IFNy (25 ng ml^-1^, StemCell Technologies), TNFα (25 ng ml^-1^, R&D Systems), IL-1β (25 ng ml^-1^, R&D Systems), LPS (100 ng ml^-1^, Fisher Scientific), diABzL (1 μM, Invivogen), cGAMP (10 μg ml^-1^, Invivogen), poly-I:C (5 μg ml^-1^, Invivogen).

### siRNA transfection

Transfection of primary iWAT adipocytes with siRNA was performed by modifying an existing protocol^58^. At day 4 of differentiation, adipocytes were trypsinized, washed and resuspended in adipocyte culture medium containing 1 μM rosiglitazone and 870 nM insulin, and for most experiments were plated at a density of 50,000 cells cm^-2^. Recipient tissue culture wells contained a 60 μl cm^-2^ solution of Opti-MEM (Gibco) with 50 nM DsiRNA duplex (Integrated DNA Technologies) and 0.5% Lipofectamine RNAiMAX (Invitrogen). For cellular respiration studies, transfected cells were plated in Seahorse XF 24-well cell culture plates using 30 μl of an Opti-MEM siRNA solution as above and a density of 18,000 cells per well. Adipocyte culture medium was changed 48 h later.

### Cellular respiration assays

Primary iWAT adipocytes were differentiated, transfected with siRNA as described above, and plated in Seahorse XF 24-well cell culture plates. At day 10 of differentiation, oxygen consumption was measured using a Seahorse XFe24 Analyzer (Agilent). Basal respiration was measured in DMEM containing 2% fatty-acid-free BSA (Roche). Cells were treated with 100 μM isoproterenol to stimulate respiration, followed by 5 μM oligomycin to inhibit coupled respiration. Maximal respiration was achieved by treatment with 100 nM valinomycin (Sigma), followed by treatment with 3 μM antimycin A (Sigma) and 3 μM rotenone (Sigma) to block mitochondrial respiration. Basal respiration were measured in 3 cycles (each cycle: 3 min mix, 2 min wait, 3 min measure), stimulated respiration in 5 cycles (1 min mix, 0 min wait, 2 min measure), uncoupled respiration in 3 cycles (2 min mix, 1 min wait, 2 min measure), maximal respiration in 3 cycles (1 min mix, 0 min wait, 2 min measure), and non-mitochondrial respiration in 2 cycles (3 min mix, 2 min wait, 3 min measure). Measurements were normalized to protein content measured using bicinchoninic acid assay (Pierce). Mitochondrial respiration was calculated by subtracting non-mitochondrial respiration from total respiration.

### Isolation of adipocytes ex vivo

Inguinal WAT was dissected, washed with PBS, and minced. It was then digested in a solution of Krebs-Ringer Bicarbonate Buffer (KRB; Sigma) containing 0.5% fatty acid-free BSA (Roche), 10 mg ml^-1^ Collagenase D (Roche), 3 U ml^-1^ Dispase II (Roche), and 5 mM CaCl_2_ for 45 min at 37 °C with shaking. The digestion solution was triturated every 10 min. After digestion, EDTA (Invitrogen) was added to a concentration of 10 mM and the cell suspension was passed through a 100 μm cell strainer. Adipocytes were floated by centrifugation at 150 g for 8 min, gently transferred to a new vessel using a wide-bore pipet, and washed with KRB. Next, a centrifugation of 150 g for 5 min was performed, after which the infranatant was removed with a syringe and the isolated adipocytes were harvested for RNA or protein analysis.

### Polysome fractionation

Inguinal WAT adipocytes were isolated by flotation as above, using 3-4 mice (6-8 iWAT lobes) per replicate sample. After isolation, adipocytes were lysed in polysome lysis buffer (20 mM HEPES-KOH pH 7.4, 100 mM KCl, 5 mM MgCl_2_, 1% Triton X-100, 2 mM DTT, 100 μg ml^-1^ cycloheximide [Sigma], 20 U ml^-1^ RNase inhibitor [Promega], and 1x Complete EDTA-free protease inhibitor cocktail [Roche]). Lysates were centrifuged 2,000 g for 5 min at 4 °C and soluble fraction recovered. Lysates were then centrifuged 15,000 g for 10 min and total RNA concentration measured in the soluble fraction by A260. A volume of lysate containing 125 ug RNA was loaded on 11 ml 15-50% sucrose gradients in 14 x 89 mm centrifuge tubes (Beckman Coulter). (Gradients were prepared in 20 mM HEPES-KOH pH 7.4, 100 mM KCl, 5 mM MgCl_2_, 2 mM DTT, 100 μg ml^-1^ cycloheximide, and 20 U ml^-1^ RNase inhibitor [Promega].) Gradients were centrifuged 36,000 rpm for 2.5 h in a SW41 Ti rotor (Beckman Coulter) with ‘slow’ acceleration and ‘max’ deceleration until brake disengagement at 2,500 rpm. Following centrifugation, 12 fractions were collected. After fraction collection, 10 pg luciferase mRNA (Promega) was spiked into each fraction as a recovery control, and RNA was isolated from each fraction using Trizol (Invitrogen) and 30 μg GlycoBlue (Thermo Fisher) as coprecipitant. Complementary DNA was prepared using a High-Capacity cDNA Reverse Transcription kit (Applied Biosystems) and mRNAs present in each fraction were determined by RT-qPCR as below.

### mRNA expression analysis

Total RNA was extracted from frozen tissue or cells using TRIzol (Invitrogen), purified with RNeasy Mini spin columns (Qiagen), and reverse transcribed using a High-Capacity cDNA Reverse Transcription kit (Applied Biosystems). The resulting cDNA was analyzed by RT-qPCR using SYBR green fluorescent dye 2x qPCR master mix (Promega) in a QuantStudio 6 Flex Real-Time PCR System (Applied Biosystems). The *Tbp* mRNA was used as a loading control, and fold change was calculated using the ΔΔCt method. Primer sequences are listed in Table S1^59^.

### RNA sequencing

Libraries were prepared using Kapa stranded mRNA Hyper Prep sample preparation kits from 500 ng of total RNA isolated using RNeasy columns (Qiagen). The finished dsDNA libraries were quantified by Qubit fluorometer (Thermo Fisher), TapeStation 2200 (Agilent), and RT-qPCR using the Kapa Biosystems library quantification kit. Uniquely indexed libraries were pooled in equimolar ratios and sequenced on an Illumina NovaSeq with paired-end 150 bp reads by the Dana-Farber Cancer Institute Molecular Biology Core Facilities.

Sequenced reads were trimmed using fastp (v0.23.2)^60^, then aligned to the mm10 reference genome assembly and quantified using STAR (v2.7.8)^61^. Differential gene expression testing was performed by DESeq2 (v.2.11.40.7)^62^, and normalized counts were used for calculation of Z-scores. Fold-change values determined by

DESeq2 were used to assemble pre-ranked gene lists for gene set enrichment analysis (GSEA) using ‘Classic’ mode for enrichment statistic calculation^63–65^. Genes with mean expression less than 5 were not included in these pre-ranked gene lists. GSEA p values are reported as familywise-error rate corrected (FWER) values. Additional gene set overrepresentation analysis was performed using non-ranked gene lists using Hallmark Gene Sets and WebGestalt^65,66^.

### Immunoblot

Tissue or cell culture samples were prepared in ice-cold cell lysis buffer (20 mM Tris pH 7.6, 150 mM NaCl, 50 mM NaF, 1 mM EDTA, 1 mM EGTA, and 1% Triton X-100, supplemented with Complete EDTA-free protease inhibitor [Roche]). Cell samples were rotated for 10 min at 4 °C to achieve lysis, whereas tissue samples were homogenized using a Polytron PT 10-35GT homogenizer (Kinematica). Lysates were sonicated for 7.5 min (30 s on, 30 s off cycles) using a Bioruptor waterbath sonicator (Diagenode), then cleared by centrifugation at 16,000 g for 15 min at 4 °C. The soluble fraction was separated from insoluble pellet and lipid and its concentration was determined by bicinchoninic acid assay (Pierce), followed by denaturation in Laemmli buffer (50 mM Tris pH 6.8, 2% SDS, 10% glycerol, 100 mM DTT, and 0.05% bromophenol blue). Proteins were resolved by SDS-PAGE in 4-12% NuPAGE Bis-Tris gels (Invitrogen) and transferred to polyvinylidene difluoride membrane with 0.45 μm pore size (Immobilon-P). Membranes were blocked with Tris-buffered saline with 0.05% Tween-20 (TBST) containing 5% dried nonfat milk (Biorad). Primary antibodies were diluted in TBST containing 5% bovine serum albumin and 0.02% NaN3, with the exception of the mouse anti-PGC1α antibody, which was diluted in TBST containing 5% dried nonfat milk (BioRad). Membranes were incubated overnight at 4 °C with primary antibody. For secondary antibody incubation, anti-rabbit HRP (Promega) or anti-mouse HRP (Promega) was diluted in TBST containing 5% dried nonfat milk. Secondary antibodies were visualized using enhanced chemiluminescence western blotting substrates (Pierce), Immobilon Crescendo HRP substrate (Millipore), or SuperSignal West Femto HRP substrate (Pierce). Densitometry was performed using ImageJ 1.52k. Primary antibodies were used to detect: PGC1α (4C1.3, Millipore); tubulin (ab4074, Abcam); TBP (44059S, Cell Signaling Technology); RPS3 (PA517214, Life Technologies).

### Mass spectrometry

Inguinal WAT adipocytes were isolated by flotation as above, using 2-3 mice (4-6 iWAT lobes) per replicate sample. After isolation, adipocytes were lysed in RIPA buffer (50 mM Tris pH 7.4, 150 mM NaCl, 1% NP-40, 0.5% sodium deoxycholate, 0.1% SDS, 2x Complete EDTA-free protease inhibitor [Roche]), and insoluble material and lipid were cleared by centrifugation at 16,000 g for 15 min at 4 °C. Protein lysates were measured using bicinchoninic acid assay (Pierce), then 25 μg of each sample was dissolved in 200 mM EPPS pH 8.5, 8 M urea, 0.1% SDS, and 1x Complete EDTA-free protease inhibitor (Roche). TCEP was added to 5 mM and protein reduction was carried out at 22 °C for 15 min, followed by alkylation by 10 mM iodoacetamide for 20 min in darkness. The reaction was quenched by 10 mM DTT for 10 min in darkness, followed by buffer exchange to 200 mM EPPS pH 8.5 using a modified Sp3 protocol^67^ (Cytiva). Protein was digested with LysC (1:33 enzyme:protein ratio) overnight at 25 °C with shaking, then trypsin was added (1:33 enzyme:protein ratio) followed by 6 hr digestion at 37 °C with shaking and then sample adjustment to a concentration of 30% acetonitrile. Peptides were labeled using 100 ug of 16-plex tandem mass tag (TMTpro) reagents (Thermo Fisher), with incubation for 60 min at 22 °C. After confirmation of labeling (>97%), excess TMTpro reagents were quenched by addition of hydroxylamine to a concentration of 0.3%. Labeled peptides were pooled and acidified with formic acid to a pH ~2 and acetonitrile concentration <2%, followed by desalting using a Sep-Pak Vac 50 mg tC18 cartridge (Waters). Peptides were eluted in 70% acetonitrile with 1% formic acid and dried by vacuum centrifugation, then resuspended in 10 mM ammonium bicarbonate pH 8 with 5% acetonitrile and fractionated into 24 fractions by basic pH reverse-phase HPLC. The fractions were dried, resuspended in 5% acetonitrile with 1% formic acid, and desalted by stage-tip, eluting in 70% acetonitrile with 1% formic acid. Samples were dried and resuspended in 5% acetonitrile with 5% formic acid. Twelve of the 24 fractions were analyzed by LC-MS/MS.

### Mass spectrometry data acquisition

Data were collected on an Orbitrap Fusion Lumos mass spectrometer (Thermo Fisher) coupled to a Proxeon EASY-nLC 1000 LC pump (Thermo Fisher). Peptides were separated using a 180-min gradient at 500 nl min^-1^ on a 30 cm column (i.d. 100 μm, Accucore, 2.6 μm, 150 Å) packed in-house. MS1 precursor scans were acquired in the orbitrap at 120 K resolution, 1e6 AGC target with a maximum of 50 ms injection time. Data dependent, “top 10” MS2 scans were acquired in the ion trap with collisional induced dissociation (CID) fragmentation. While setting varied from instrument to instrument and with instrument performance, generally the following was used for MS2 scans: NCE 35%, 1.2e4 AGC target, maximum injection time 50 ms, isolation window 0.5 Da. Orbiter, an on-line real-time search algorithm, was used to trigger MS3 quantification scans^68^. MS3 scans were acquired in the orbitrap using varying settings based on instrument performance: 50,000 resolution, AGC of 2.5 × 10^5^, injection time of 200 ms, HCD collision energy of 65%. Protein-level closeout was typically set to two peptides per protein per fraction.

### Mass spectrometry data analysis

Raw files were first converted to mzXML, and monoisotopic peaks were re-assigned using Monocle^69^. Database searching included all mouse entries from Uniprot (downloaded in July, 2014). The database was concatenated with one composed of all protein sequences in the reversed order. Sequences of common contaminant proteins (e.g., trypsin, keratins, etc.) were appended as well. Searches were performed using the comet search algorithm. Searches were performed using a 50 ppm precursor ion tolerance and 1.0005 Da fragment ion tolerance. TMTpro on lysine residues and peptide N termini (+304.2071 Da) and carbamidomethylation of cysteine residues (+57.0215 Da) were set as static modifications, while oxidation of methionine residues (+15.9949 Da) was set as a variable modification.

Peptide-spectrum matches (PSMs) were adjusted to a 1% false discovery rate (FDR)^70^. PSM filtering was performed using linear discriminant analysis (LDA) as described previously^71^, while considering the following parameters: comet log expect, different sequence delta comet log expect (percent difference between the first hit and the next hit with a different peptide sequence), missed cleavages, peptide length, charge state, precursor mass accuracy, and fraction of ions matched. Each run was filtered separately. Protein-level FDR was subsequently estimated at a data set level. For each protein across all samples, the posterior probabilities reported by the LDA model for each peptide were multiplied to give a protein-level probability estimate. Using the Picked FDR method^72^, proteins were filtered to the target 1% FDR level.

For reporter ion quantification, a 0.003 Da window around the theoretical *m/z* of each reporter ion was scanned, and the most intense *m/z* was used. Reporter ion intensities were adjusted to correct for the isotopic impurities of the different TMTpro reagents according to manufacturer specifications. Peptides were filtered to include only those with a summed signal-to-noise (SN) of 160 or greater across all channels. For each protein, the filtered peptide TMTpro SN values were summed to generate protein quantification. Fold-change values were used to generate a pre-ranked protein list for gene set enrichment analysis using the same methodology as employed for RNA-seq analyses.

### Body composition measurement

Fat and lean mass composition of living mice was assessed using a 3-in-1 Echo MRI Composition analyzer (Echo Medical Systems).

### Diet-induced obesity

Age-matched littermates and group housing were used for high-fat feeding experiments. At 8 weeks of age, mice were given 60 kcal% fat rodent diet D12492 (Research Diets, Inc.) ad libitum. Mouse weight and consumed food weight were assessed once per week.

### Intraperitoneal glucose tolerance test

Mice were fasted overnight, after which their weight and fasting blood glucose values were measured using tail nick blood and a OneTouch UltraMini glucose meter. Glucose was administered by intraperitoneal injection at a dose of 1 g kg^-1^, followed by blood glucose measurements over the next 3 h.

### Indirect calorimetry

Mice were individually housed in Promethion metabolic cages (Sable Systems International) at 22 °C with a 12 h light-dark cycle. After at least 2 d acclimatization to the cages, mice were injected intraperitoneally at Zeitgeber time ZT4 with 100 μl PBS and returned to their cages for measurement. The next day, the same volume of the β3-adrenergic receptor agonist CL-316,243 was administered intraperitoneally at a dose of 1 mg kg^-1^. Energy expenditure measurements were made every 3 min thereafter for 2.5 h using Sable Systems data acquisition software. Data were processed using Sable Systems MacroInterpreter software using One-Click Macro. Individual measurements were not normalized to body weight, and body weight did not significantly differ between the experimental groups.

### Statistical analyses

Replicate numbers are described in figure legends. For cellular assays, n corresponds to the number of experimental replicates (e.g., independent transfections). For animal assays or tissue extracted from animals, n corresponds to the number of mice used per genotype or condition. Sample sizes were determined on the basis of previous experiments using similar methodologies. Unless otherwise stated, data are presented as mean and error bars indicate standard error. Graphing and statistical analyses, including two-tailed Student’s *t*-test, one-way ANOVA, and Fisher’s LSD, were performed using Prism (GraphPad).

## SUPPLEMENTAL FIGURE LEGENDS

**Figure S1.**
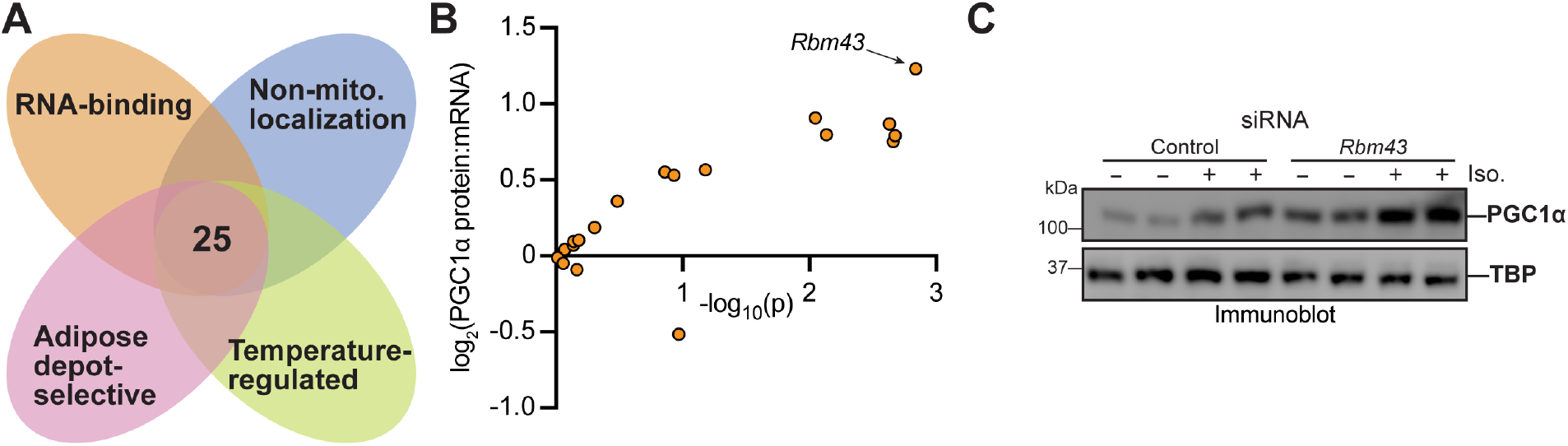
Candidate post-transcriptional regulators of oxidative metabolism in adipocytes. **(A)** Criteria for identification of RNA-binding proteins with potential roles in regulation of adipocyte oxidative metabolism. Adipocyte-specific TRAP data were used for gene expression analyses (NCBI GEO: GSE108077). **(B)** Candidate proteins were knocked down using siRNA in primary iWAT adipocytes. For each successful knockdown, the resulting change in PGC1α protein:mRNA ratio is plotted versus its statistical significance. n = 3 (RNA), 4 (protein). **(C)** Immunoblot of PGC1α protein levels upon *Rbm43* knockdown in primary iWAT adipocytes. Samples were treated as indicated with 100 nM adrenergic agonist isoproterenol (3 h) to induce *Ppargc1a* transcription.

**Figure S2.**
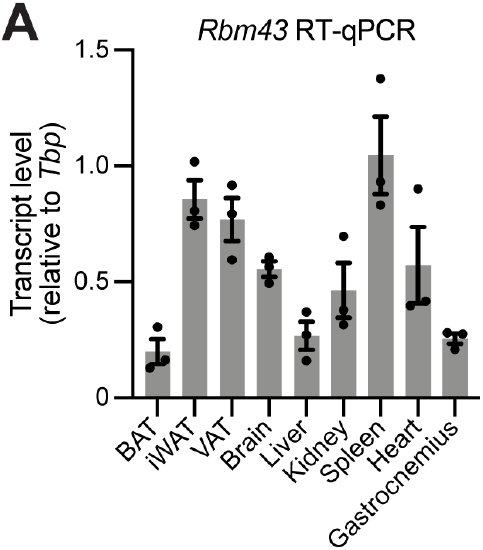
Expression of *Rbm43* in mouse tissues. **(A)** *Rbm43* mRNA was assessed by RT-qPCR in indicated tissues of male mice housed at 22 °C. n=3.

**Figure S3.**
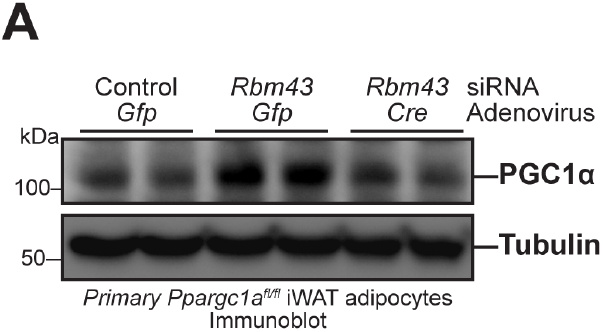
Normalization of PGC1α in *Rbm43* knockdown cells via Cre-mediated *Ppargc1a^fl/fl^* deletion. **(A)** Immunoblot of PGC1α upon *Rbm43* knockdown in *Ppargc1a^fl/fl^* primary iWAT adipocytes in the presence of adenoviral *Cre* or *Gfp*.

**Figure S4.**
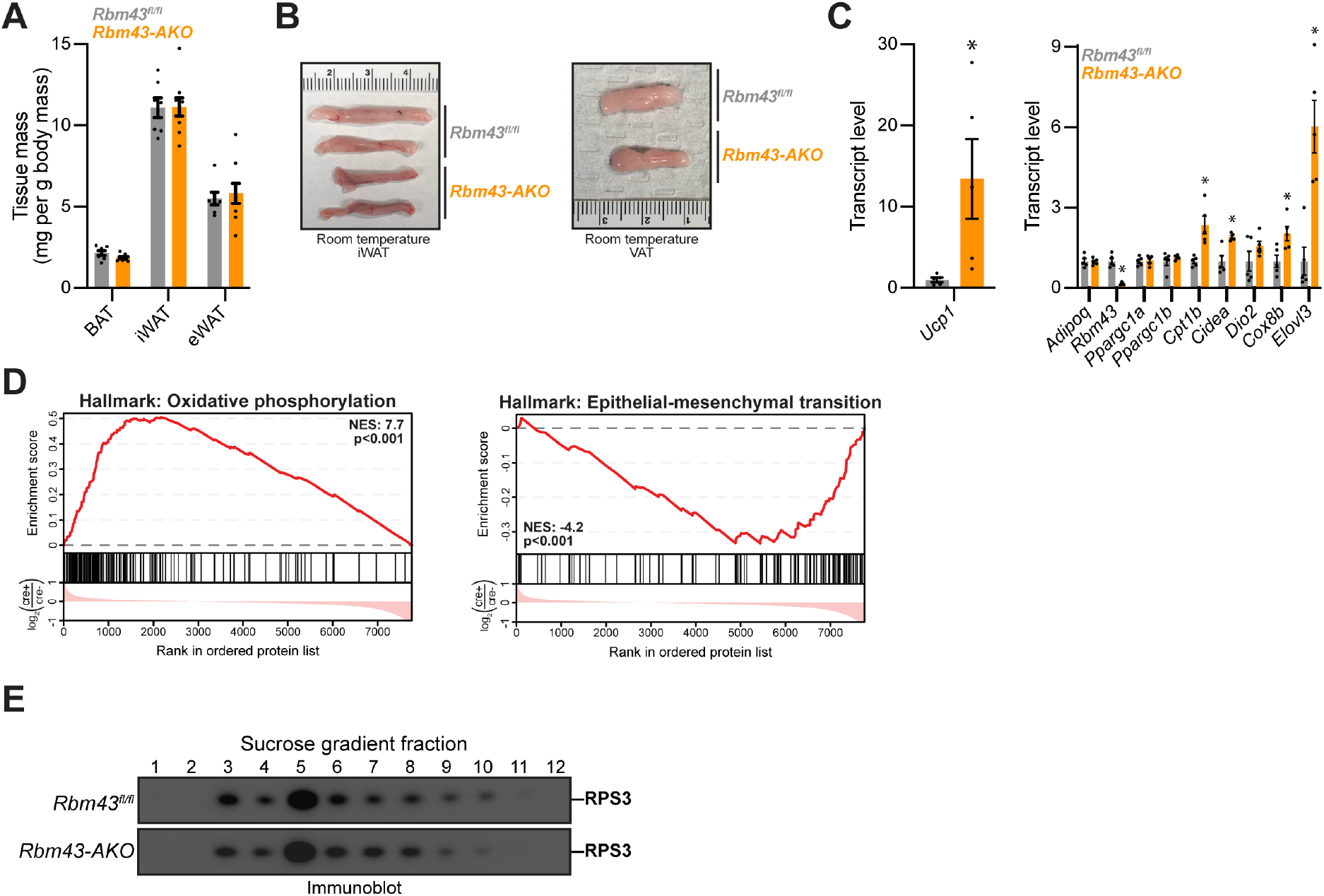
Adipocyte-selective knockout of *Rbm43*. **(A)** Mass of fat depots dissected from female *Rbm43^fl/fl^* and *Rbm43-AKO* mice fed a chow diet. n=7-9. **(B)** Gross appearance of fat depots dissected from *Rbm43^fl/fl^* and *Rbm43-AKO* mice. **(C)** RT-qPCR assessing *Rbm43^fl/fl^* and *Rbm43-AKO* iWAT adipocytes. n=5 flotation experiments, each including 2-3 mice. *p<0.05. **(D)** Gene set enrichment plots assessing enrichment of oxidative phosphorylation (left) and epithelial-mesenchymal transition (right) gene sets in *Rbm43-AKO* iWAT proteomics data. **(E)** Immunoblot of Rps3 in iWAT adipocyte *Rbm43^fl/fl^* and *Rbm43-AKO* polysome profile fractions.

**Figure S5.**
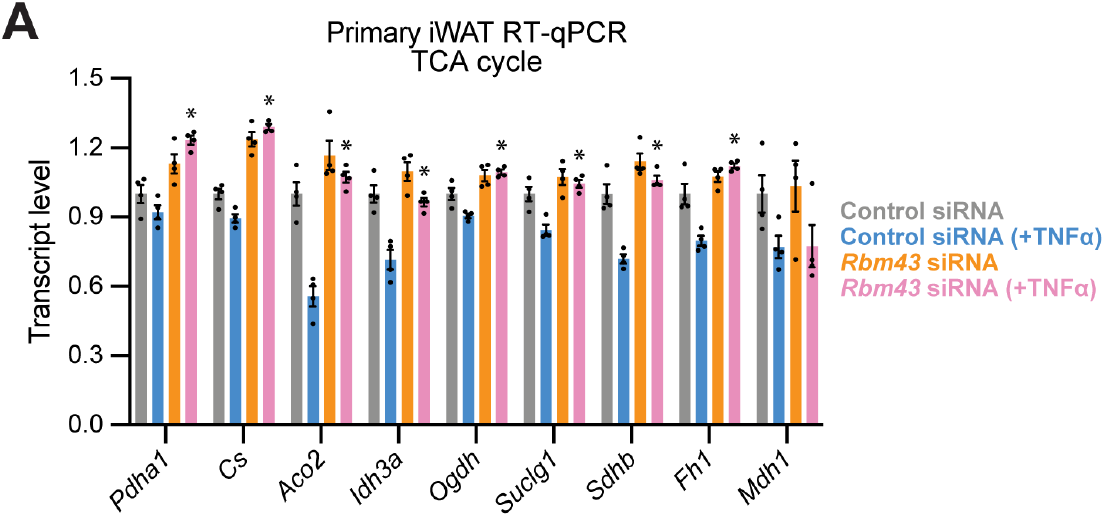
RBM43 loss confers protection from cytokine-induced impairment in mitochondrial gene expression. **(A)** RT-qPCR assessing expression of TCA cycle components in primary iWAT adipocytes after 24 h treatment with TNFα, in the presence or absence of *Rbm43* knockdown. n=4. *p<0.005 vs control siRNA (+TNFα).

