## Supplemental Table S1 for "RBM43 links adipose inflammation and energy expenditure through translational regulation of PGC1α"

**Table S1. Primers used for RT-qPCR of mouse genes**

| Gene | Primers |
|---|---|
| *Tbp* | GAAGCTGCGGTACAATTCCAG |
| | CCCCTTGTACCCTTCACCAAT |
| *Fabp4* | AAGGTGAAGAGCATCATAACCCT |
| | TCACGCCTTTCATAACACATTCC |
| *Rbm43* | GCATCAGCGTGGAAGGTCAA |
| | CCCTCGTCTTGGAAGTAGCAC |
| *Rbm43* | ATCAGCGTGGAAGGTCAAAG |
| | GTCCACCCTCGTCTTGGAAG |
| *Ppargc1a* | GGTTGGTGAGGACCAGCC |
| | AATCCACCCAGAAAGCTGTCT |
| *Cpt1b* | GCACACCAGGCAGTAGCTTT |
| | CAGGAGTTGATTCCAGACAGGTA |
| *Cidea* | TGACATTCATGGGATTGCAGAC |
| | GGCCAGTTGTGATGACTAAGAC |
| *Ucp1* | AGGCTTCCAGTACCATTAGGT |
| | CTGAGTGAGGCAAAGCTGATTT |
| *Dio2* | AATTATGCCTCGGAGAAGACCG |
| | GGCAGTTGCCTAGTGAAAGGT |
| *Cox8b* | TGTGGGGATCTCAGCCATAGT |
| | AGTGGGCTAAGACCCATCCTG |
| *Cs* | GGACAATTTTCCAACCAATCTGC |
| | TCGGTTCATTCCCTCTGCATA |
| *Cycs* | GCAAGCATAAGACTGGACCAAA |
| | TTGTTGGCATCTGTGTAAGAGAATC |
| *Retn* | AAGAACCTTTCATTTCCCCTCCT |
| | GTCCAGCAATTTAAGCCAATGTT |
| *Ppargc1b* | TGCGGAGACACAGATGAAGA |
| | GGCTTGTATGGAGGTGTGGT |
| *Elovl3* | TTCTCACGCGGGTTAAAAATGG |
| | GAGCAACAGATAGACGACCAC |
| *Pdha1* | GAAATGTGACCTTCATCGGCT |
| | TGATCCGCCTTTAGCTCCATC |
| *Aco2* | ATCGAGCGGGGAAAGACATAC |
| | TGATGGTACAGCCACCTTAGG |
| *Idh3a* | GTTGCTTCGTAAGACATTTGACC |
| | TCTCTCGGATGGTGACGATATT |
| *Ogdh* | AGGGCATATCAGATACGAGGG |
| | CTGTGGATGAGATAATGTCAGCG |
| *Suclg1* | TGGGATACGACACGGGTCTTA |
| | CAGAAGCCGTTGCTCCTGTT |
| *Sdhb* | AATTTGCCATTTACCGATGGGA |
| | AGCATCCAACACCATAGGTCC |
| *Fh1* | GAATGGCAAGCCAAAATTCCTT |
| | CGTTCTGTAGCACCTCCAATCTT |
| *Mdh1* | TTCTGGACGGTGTCCTGATG |
| | TTTCACATTGGCTTTCAGTAGGT |
